# The gut microbiota metabolite isovalerate enhances the epithelial barrier function in cell monolayers derived from porcine ileum organoids

**DOI:** 10.1101/2025.06.04.657803

**Authors:** Martin Beaumont, Cláudia M. Vicente, Corinne Lencina, Elisabeth Jones, Stéphanie Lecuelle, Tristan Chalvon-Demersay

## Abstract

The gut microbiota produces numerous metabolites that influence the epithelial barrier function. Bacterial catabolism of amino acids produces a wide variety of metabolites whose effects on the intestinal epithelium remain to be fully identified. In this study, we investigated the effects of amino acid derived metabolites (isovalerate, isobutyrate, 2-methylbutyrate, 5-aminovalerate, cadaverine, putrescine, and tryptamine) in cell monolayers derived from porcine ileum organoids. Our results show that the leucine-derived branched-chain fatty acid (BCFA) isovalerate improved the epithelial barrier function, as assessed by transepithelial electrical resistance measurement and permeability assay. Isovalerate upregulated the expression of genes involved in innate immunity, markers of absorptive cells and enteroendocrine cells while reducing the expression of the stem cells and mucus related genes. Most of the effects of isovalerate on epithelial cells were also observed with the bacterial metabolite butyrate, an inhibitor of the epigenetic enzymes histone deacetylases (HDAC). Furthermore, the structurally unrelated HDAC inhibitor trichostatin A improved epithelial barrier function and upregulated *SLPI* gene expression, as observed with isovalerate and butyrate. Isovalerate also upregulated the gene expression of antioxidant enzymes and this effect was not observed with butyrate. Interestingly, the other two BCFAs isobutyrate and 2-methylbutyrate did not replicate the effects of isovalerate, suggesting that the carbon chain structure of isovalerate mediates its effect. In contrast, we found that all three BCFAs were able to cross the epithelial cell monolayer derived from porcine ileum organoids from the apical to the basal side. Overall, our *in vitro* results suggest that targeting the bacterial production of isovalerate may be useful to promote gut health. In this perspective, we performed an *in silico* analysis that identified species belonging to dominant gut microbiota genera such as *Prevotella, Blautia, Christensenella, Clostridium,* and *Ruminococcus,* as potential producers of BCFAs through the PorA enzymatic pathway.

## Introduction

The intestinal epithelium contributes to digestion and nutrient absorption while forming a physiochemical and immunological barrier in the gut against microorganisms, toxins and food antigens (Peterson and Artis 2014). These functions of the intestinal epithelium rely on diverse epithelial cell types including absorptive cells, BEST4^+^ cells, goblet cells, Paneth cells and enteroendocrine cells, all derived from rapidly cycling stem cells (Elmentaite et al. 2021; Malonga et al. 2024). The barrier function of the intestinal epithelium involves the formation of tight junctions, the sensing of microbes and the release of mucus and antimicrobial peptides (Peterson and Artis 2014). The gut microbiota is a key regulator of the epithelial barrier function, notably through the release in the lumen of bacterial metabolites able to diffuse through the mucus layer and reach epithelial cells (Ghosh et al. 2021). The identification of bacterial metabolites able to regulate the epithelial barrier function and the understanding of their mode of action could lead to the development of innovative tools to promote gut health.

Short-chain fatty acids are the main end products of the fermentation of carbohydrates by the gut microbiota (Oliphant and Allen-Vercoe 2019; van der Hee and Wells 2021). Among them, butyrate is well known for its protective role for the epithelial barrier function (Peng et al. 2009; Pearce et al. 2020; Alberge et al. 2024). The gut microbiota also produces a wide diversity of metabolites from amino acids but their effects on the intestinal barrier function remain to be fully elucidated (Portune et al. 2016; Oliphant and Allen-Vercoe 2019; Beaumont et al. 2022). These amino acids can either be derived from incompletely digested dietary proteins, from host secretions, or from bacterial proteins. Reductive deamination of the branched-chain amino acids leucine, valine and isoleucine produces the branched-chain fatty acids (BCFAs) isovalerate, isobutyrate and 2-methylbutyrate, respectively (Allison 1978; Van den Abbeele et al. 2022). BCFAs are recognized as reliable markers of amino acid fermentation in the gut (Gilbert et al. 2018). The few studies that evaluated the effects of BCFAs on the intestinal epithelium suggested their protective effect for the barrier function (Boudry et al. 2013; Ezzine et al. 2022; Wang et al. 2023). Microbial catabolism of arginine releases the polyamine putrescine, which was shown to modulate mitochondrial metabolism and barrier function in epithelial cells (Bekebrede et al. 2020). Decarboxylation of lysine releases the polyamine cadaverine and 5-aminovalerate whose effects on the epithelium have not been characterized (Portune et al. 2016; Oliphant and Allen-Vercoe 2019). Decarboxylation of tryptophan by the gut microbiota releases numerous metabolites including tryptamine, which is an indole containing monoamine (Williams et al. 2014; Roager and Licht 2018). Tryptamine is known as a neurotransmitter, accelerates gut transit through increased colonic secretion, increases mucus secretion and protects mice from colitis (Williams et al. 2014; Bhattarai et al. 2018, 2020). Overall, amino acid derived metabolites may play a key role in the microbial regulation of the gut barrier function but a more detailed characterization of their effects on the intestinal epithelium is required.

The aim of our study was to investigate the effects of gut microbiota metabolites derived from amino acids by using an *in vitro* organoid model that recapitulates the cellular diversity and functions of the intestinal epithelium. We used organoids derived from the porcine intestine as this species is considered as an excellent model for biomedical research (Ziegler et al. 2016). Furthermore, protein fermentation has been linked with gut health in pigs (Gilbert et al. 2018) and the concentration of putrescine, cadaverine, and 5-aminovalerate is reduced in the gut of piglets after weaning, which is a key step in intestinal development (Beaumont et al. 2021). Porcine intestinal organoids retain characteristics of their digestive segment of origin (Mussard et al. 2022). Therefore, in this study we used organoids derived from the pig ileum as a likely anatomical site for the production of gut microbiota metabolites derived from amino acids. Indeed, amino acids are absorbed in the small intestine where the microbial density is the highest in the distal ileum. In order to expose epithelial cells to amino acid derived metabolites through their apical side, we treated porcine ileal organoid cells cultured in a 2D monolayer format (Mussard et al. 2023).

## Materials and methods

### Culture of porcine ileum organoids

Ileum organoids derived from suckling piglets (21-day-old) were obtained from our in-house biobank and cultured as described before (Mussard et al. 2022, 2023; Alberge et al. 2024). Briefly, frozen ileum crypts or organoids (passages 1 to 5) kept in liquid nitrogen were thawed at 37°C, centrifuged (500 g, 4°C, 5 min) and seeded in Matrigel (Corning, cat#354234) in a pre-warmed 48-well plate (25 µL/well). Organoid culture medium containing IntestiCult Organoid Growth Medium (Human) (StemCell Technologies, cat#6010) supplemented with 1% Penicillin-Streptomycin (Sigma, cat#P4333) and 100 µg/mL Primocin (InvivoGen, cat#ant-pm-05) was added (250 µL/well). Organoids were cultured at 37°C with 5% CO_2_. Five to ten days after seeding, organoids in Matrigel were washed in PBS (ThermoFischerScientific, cat#10010015) and homogenized by pipetting in warm TrypLE (ThermoFischerScientific, cat#12605-010) before incubation for 15 min at 37°C. Digestion was stopped by adding cold complete DMEM (DMEMc) containing DMEM (ThermoFischerScientific, cat#31966047) supplemented with 10% fetal bovine serum (FBS, ThermoFischerScientific, cat#10270-106) and 1% Penicillin-Streptomycin. Cells were centrifuged (500 g, 4°C, 5 min) and counted using a Countess 3 Automated Cell Counter (ThermoFischerScientific, cat#16842556). Organoid cells were seeded in Matrigel:DMEMc (v/v: 2:1) in pre-warmed 24-well plates (3 000 cells/50 µL/well) and organoid culture medium was added (500 µL/well) and replaced every 2-3 days. Organoids (passages 1 to 6) were used to seed cell monolayers 7 days after seeding.

### Culture of cell monolayers derived from porcine ileum organoids

Cell culture inserts for 24-well plates (Corning, cat#353095) were coated with 50 µg/mL Collagen type IV from human placenta (Sigma, cat#C5533) for 2 h at 37°C (150 µL/well). The coating solution was removed and the inserts were dried for 10 min by opening the plate lid under the cell culture cabinet. Organoids were dissociated and cells were counted and centrifuged as described above. Cells were resuspended in organoid culture medium supplemented with 20% FBS and 10 µM Y27632 (ATCC, cat#ACS-3030) before seeding in inserts (2.5 10^5^ cells/insert). The same medium was used at the basal side. Cell culture inserts were placed in a 24-well plate (apical volume: 200 µL, basal volume: 500 µL) or in a CellZscope+ system (nanoAnalytics) used for automatic measurement of transepithelial electrical resistance (TEER, apical volume: 400 µL, basal volume: 770 µL). Cells were incubated at 37°C, 5% CO_2_. All experiments were repeated with ileum organoid cell monolayers derived from at least three 21-day old suckling piglets.

### Treatments of cell monolayers derived from porcine ileum organoids

#### Experiment 1 (screening of the effects of bacterial metabolites)

One day after seeding, the apical medium was replaced by organoid culture medium supplemented with bacterial metabolites (1 mM) or PBS (negative control). Stock solution of the following bacterial metabolites (100 mM, all from Sigma) were prepared in PBS: butyrate (cat#B5887), cadaverine (cat#C8561-G), 5-aminovalerate (cat#194336-5G), putrescine (cat#P5780-5G), tryptamine (cat#246557-5G), isobutyrate (cat#I1754-100ML), isovalerate (cat#129542-100ML), and 2-methylbutyrate (cat#W269514-SAMPLE-K). After 24h of treatment, the transepithelial electrical resistance (TEER) was measured with an EVOM3 volt/ohm meter (World Precision Instruments). For gene expression analysis, cells were lyzed in 300 µL TriReagent (Ozyme, cat# ZR2050-1-200) and kept at -80°C until RNA purification.

#### Experiment 2 (dose-response effects of butyrate and branched-chain fatty acids)

Two or three days after seeding, the apical medium was replaced by PBS supplemented with butyrate, isovalerate, isobutyrate or 2-methylbutyrate (1, 3 or 5 mM) or PBS only (negative control). The TEER was measured with a CellZscope+ system. After 48h of treatment, cells were lyzed in 300 µL TriReagent and kept at -80°C until RNA purification. Apical and basal media were collected and stored at -80°C until metabolomics analyses.

#### Experiment 3 (mode of action of butyrate and isovalerate)

Two days after seeding, the apical medium was replaced by PBS supplemented or not with butyrate 5 mM or isovalerate 5 mM or the histone deacetylase inhibitor Trichostatin A 1 µM (TSA, Sigma, cat#T1952-200UL) or the PPARγ inhibitor GW9662 10 µM (Sigma, cat# M6191-5MG) or butyrate 5 mM + GW9662 10 µM or isovalerate 5 mM + GW9662 10 µM. All cells were treated with a final concentration of 0.1% DMSO (vehicle of TSA and GW9662). The TEER was measured with a CellZscope+ system. After 48h of treatment, FITC-dextran 4 kDa (Sigma, cat#FD4-100MG) prepared in warm HBSS (Thermo Fisher Scientific, cat#15266355) was added at the apical side (2.2 mg/mL, 200 µL) of the inserts before transfer to a new 24-well plate. Warm HBSS was added at the basal side (500 µL). After incubation (2h, 37°C), fluorescence (Excitation: 495 nm, Emission: 530 nm) of the basal medium was quantified with a multimode plate reader Infinite 200 PRO (Tecan). Cells were then washed twice in PBS before lysis in 300 µL TriReagent and kept at -80°C until RNA purification.

### Gene expression analysis

RNA was purified by using the Direct-zol RNA Microprep kit (Zymo Research, cat#R2062), following the manufacturer instructions. RNA was eluted in 15 µL RNAse-free water and quantified with a NanoDrop 8000 spectrophotometer (Thermo Fisher Scientific). RNA (300 or 500 ng depending on the experiment) were reverse transcribed to cDNA by using GoScript Reverse Transcription Mix, Random primer (Promega, cat#A2801), following the manufacturer instructions. Gene expression was analyzed by real-time qPCR using QuantStudio 6 Flex Real-Time PCR System (Thermofisher) or Biomark microfluidic system using 96.96 Dynamic Arrays IFC for gene expression (Fluidigm) according to the manufacturer recommendations. The sequences of the primers used are presented in supplementary table 1. Data were normalized to the stably expressed gene HPRT1 (experiment 1) or RPL32 (experiment 2) or GAPDH (experiment 3) and analyzed with the 2^−ΔCt^ method.

### Metabolomics

Culture media (apical and basal) were centrifuged (18 000 g, 10 min, 4°C). The supernatant was collected and 50 µL was mixed with 600 µL of phosphate buffer pH 7 prepared in D_2_O and containing the internal standard TSP (1 mM). Samples were transferred in 5-mm nuclear magnetic resonance (NMR) tubes. The spectra were acquired at 300 K using the Carr-Purcell-Meiboom-Gill (CPMG) spin-echo pulse sequence with pre-saturation on a AVANCE III HD NMR spectrometer operating at 600.13 MHz for ^1^H resonance frequency using a 5 mm inverse detection CryoProbe (Bruker) at the metabolomics platform MetaToul-AXIOM (INRAE, Toulouse, France) as described before (Beaumont et al. 2020). Baseline correction, bucketing (0.01 ppm) and normalization by the total spectrum area was performed with the ASICS package (Lefort et al. 2019). The normalized intensity of the buckets corresponding to the following chemical shifts were used for metabolite quantification: 0.86 ppm for 2-methylbutyrate, 0.89 ppm for butyrate, 0.92 ppm for isovalerate, and 1.07 ppm for isobutyrate. Metabolite identifications were confirmed by comparison of sample spectra with the spectra of pure compounds obtained in the same buffer and with the same NMR spectrometer.

### Statistical analyses

Statistical analyses were performed in R (version 4.2.0). All data were log normalized before analysis with linear mixed models (R packages car, lme4, emmeans) (Bates et al. 2015; Fox et al. 2023; Lenth et al. 2024). The fixed effect was the treatment and the random effect was the pig from which organoids were derived. The mean of each group was compared to the mean of the control group (Dunnett multiple comparison procedure). For graphical representations, all results were expressed relatively to the value measured in the control sample from the same pig.

### Homologous gene cluster analyses

The search for grouped homologous genes was performed using BLAST (Camacho et al. 2009) and cblaster through the Comparative Gene Cluster Analysis Tool (CAGECAT) (Gilchrist et al. 2021). The locus pyruvate:ferredoxin oxidoreductase (POR) in *Clostridium sporogenes* consisting of *porA* and neighboring genes (∼3 kb) was used as query since this locus was previously shown to be involved in BCFA production (Guo et al. 2019). The maximum intergenic gap was set at 20 000 bp and the maximum intermediate gene distance set at 5 000 bp. The clinker tool was used for gene cluster comparison and visualization (Gilchrist and Chooi 2021).

## Results

### Screening of amino acid derived bacterial metabolites reveals that isovalerate alters gene expression in intestinal epithelial cells

We first performed a screening experiment to evaluate the effects of gut microbiota metabolites derived from amino acids (isovalerate, isobutyrate, 2-methylbutyrate, cadaverine, 5-aminovalerate, putrescine and tryptamine) on the intestinal epithelium (Figure 1A). Butyrate was used as a positive control known to influence epithelial functions. Cell monolayers derived from porcine ileum organoids were treated at the apical side with bacterial metabolites at a single concentration of 1 mM for 24h (Figure 1B). The TEER was similar to the control group in all conditions, indicating that epithelial integrity remained intact after incubation with the bacterial metabolites (Figure 2A). Accordingly, the gene expression of proteins involved in epithelial cell junctions remained unchanged (Figure 2B). In contrast, butyrate and isovalerate strongly increased the expression of the antimicrobial protein SLPI (Figure 2C). The expression of genes encoding proteins involved in redox balance remained stable under all experimental conditions (Figure 2D). Butyrate decreased the gene expression of one subunit of the NF-κB transcription factor (NFKB1) and of the apoptosis marker BAX (Figure 2E), whereas it increased the expression of the cytokine CXCL8 (Figure 2F). In contrast with butyrate, tryptamine increased the gene expression of BAX (Figure 2E).

**Figure 1:**
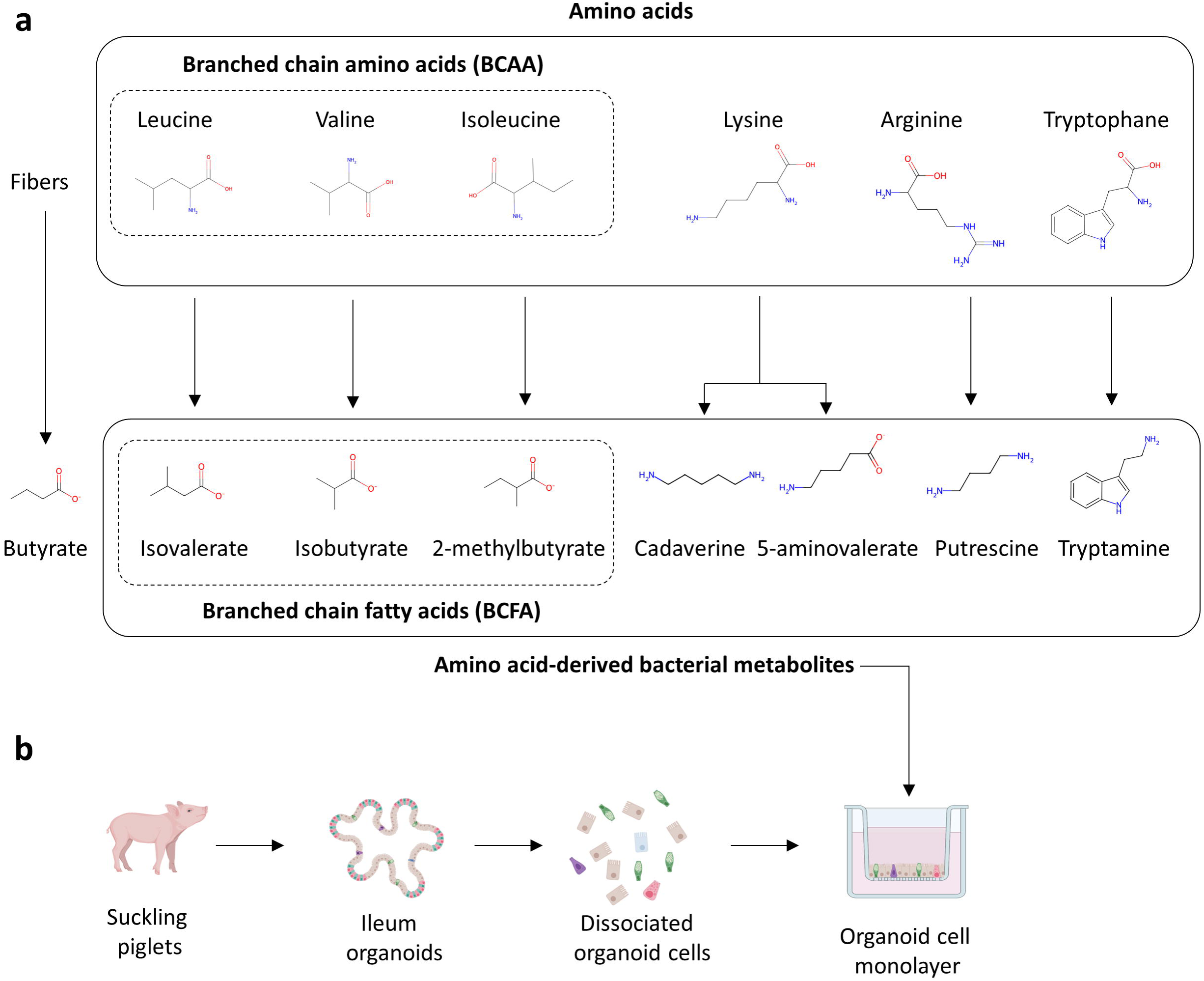
Schematic representation of the experimental design. a) The molecular structure of amino acid derived metabolites and their precursor amino acids are shown. The structure of butyrate is shown as this metabolite, which is mainly derived from carbohydrates, was used as a positive control in the experiments. b) Schematic representation of the experimental protocol used to test the effects of amino acid derived metabolites in cell monolayers derived from porcine ileum organoids.

**Figure 2:**
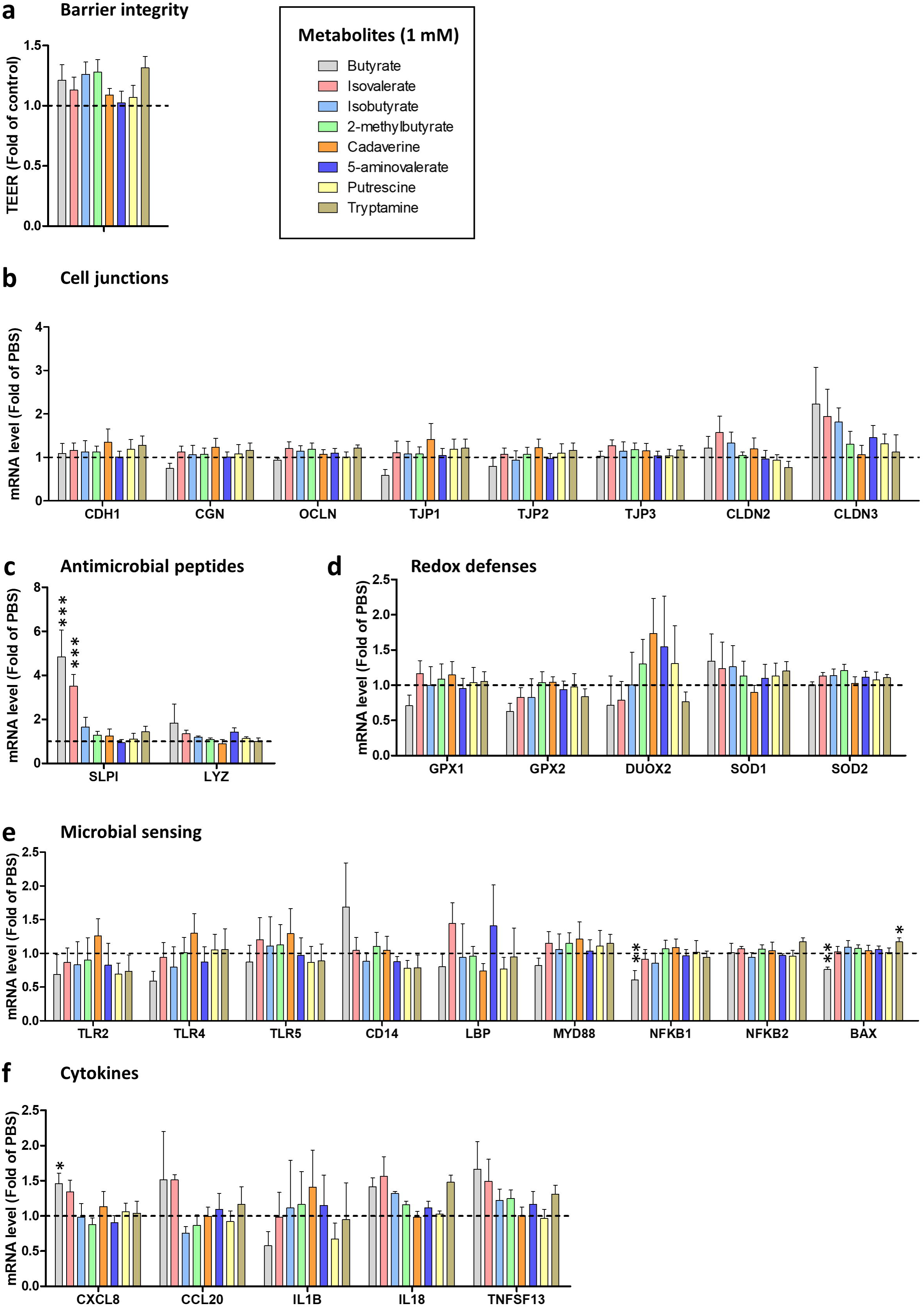
Effects of butyrate and amino acid derived metabolites on the epithelial barrier function. Cell monolayers derived from porcine ileum organoids were treated for 24h with bacterial metabolites (butyrate, isovalerate, isobutyrate, 2-methylbutyrate, cadaverine, 5-aminovalerate, putrescine, tryptamine) at 1 mM or with vehicle (PBS, negative control). a) Transepithelial electrical resistance (TEER) shown as means and SEM, n=7/group. Data are expressed relative to the control condition (PBS), represented by the dotted line (y=1). b - f) Relative gene expressions are represented as means and SEM, n>3/group. Data are expressed relative to the control condition (PBS), represented by the dotted line (y=1). *: P<0.05 versus control group, **: P<0.01 versus control group, ***: P<0.001 versus control group. b) Genes encoding proteins involved in cell junctions. c) Genes encoding antimicrobial peptides. d) Genes encoding for proteins involved in redox defenses. e) Genes encoding proteins involved in microbial sensing. f) Genes encoding cytokines.

Butyrate decreased the gene expression of the epithelial progenitor marker genes MKI67, PCNA, and CFTR, whereas it increased the expression of CDX2 (Figure 3A). Butyrate and isovalerate increased the expression of the fatty acid binding proteins FABP1 and FABP6 (Figure 3B). Tryptamine also increased the expression of FABP6 (Figure 3B). Butyrate also upregulated the expression of other markers of absorptive cell differentiation NHE3, CA2 and VIL1 (Figure 3B). Tryptamine increased the gene expression of the arginine degrading enzyme ARG2 (Figure 3C). Butyrate and isovalerate similarly enhanced the expression of the enteroendocrine marker gene CHGA (Figure 3D). The expression of genes expressed by goblet cells and involved in glycocalyx formation remained unchanged after treatment with metabolites (Figure 3E). Overall, the screening experiments revealed that isovalerate and tryptamine induce changes in gene expression in the intestinal epithelium.

**Figure 3:**
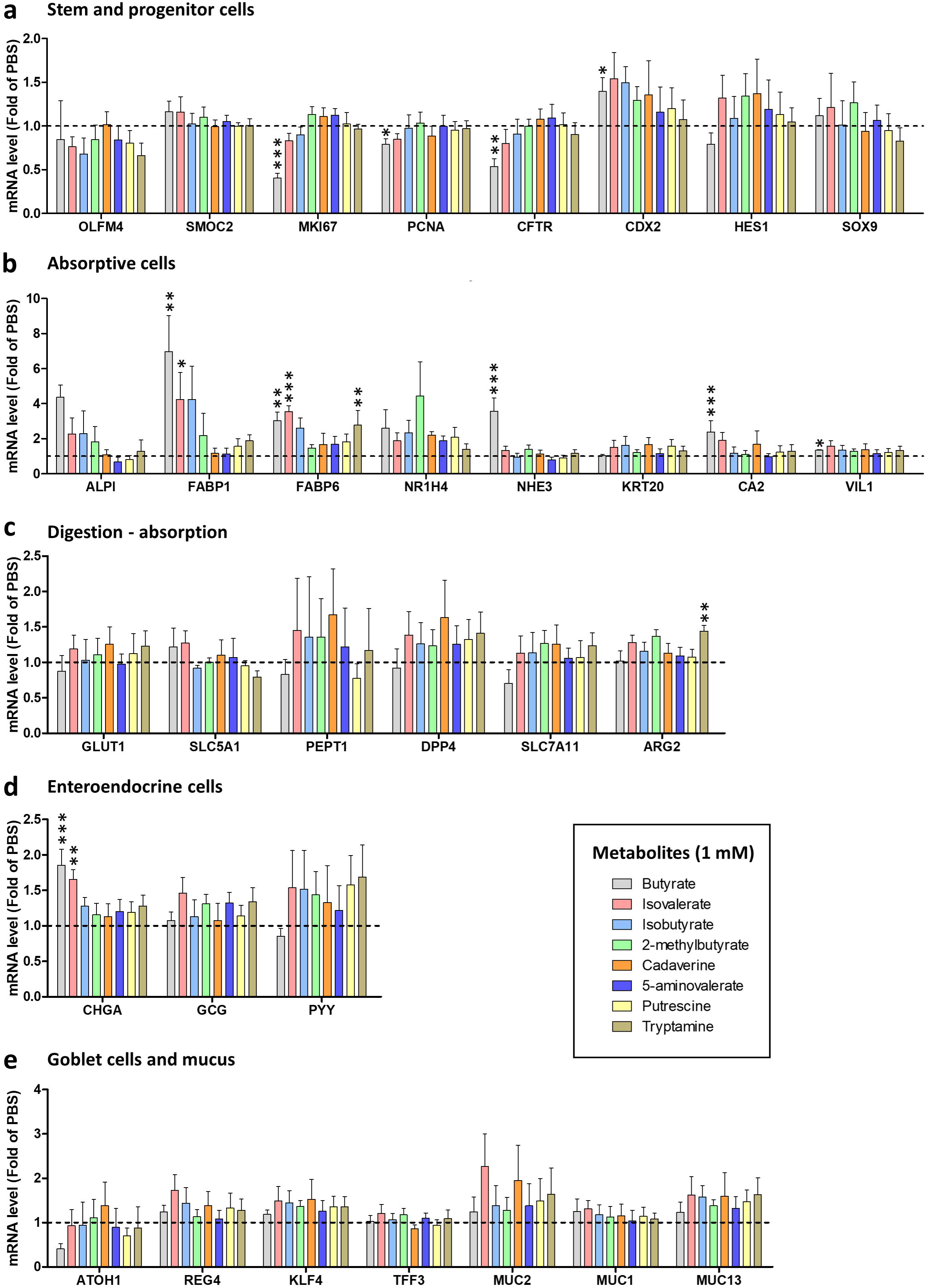
Effects of butyrate and amino acid derived metabolites on epithelial cell proliferation and differentiation. Cell monolayers derived from porcine ileum organoids were treated for 24h with bacterial metabolites (butyrate, isovalerate, isobutyrate, 2-methylbutyrate, cadaverine, 5-aminovalerate, putrescine, tryptamine) at 1 mM or with vehicle (PBS, negative control). a - e) Relative gene expressions are represented as means and SEM, n>3/group. Data are expressed relative to the control condition (PBS), represented by the dotted line (y=1). *: P<0.05 versus control group, **: P<0.01 versus control group, ***: P<0.001 versus control group. a) Marker genes for stem and progenitor cells. b) Marker genes for absorptive cells. c) Genes involved in epithelial digestion and nutrient transport. d) Marker genes for enteroendocrine cells. e) Marker genes for goblet cells and mucus.

### Dose-response experiments show that isovalerate enhances the epithelial barrier function

Following the results of the screening experiment showing that isovalerate shared some of the effects of butyrate on epithelial cells, we decided to investigate in more detail the effects of BCFAs (isovalerate, isobutyrate and 2-methylbutyrate) on the intestinal epithelium. Cell monolayers derived from porcine ileum organoids were treated at the apical side with butyrate or BCFAs at three concentrations (1, 3 or 5 mM) for 48h. Butyrate (3 and 5 mM) and isovalerate (3 and 5 mM) increased the epithelial barrier integrity from 12h of treatments, as indicated by TEER measurement (Figure 4A). The lowest concentration of butyrate (1 mM) also increased TEER from 24h of treatment. The two other BCFAs isobutyrate and 2-methylbuyrate had no effect on the TEER. Butyrate (3 and 5 mM) reduced the gene expression of the pore-forming claudin CLDN2, whereas it increased the expression of the adherens junction protein E-cadherin encoded by CDH1 (Figures 4B and C).

**Figure 4:**
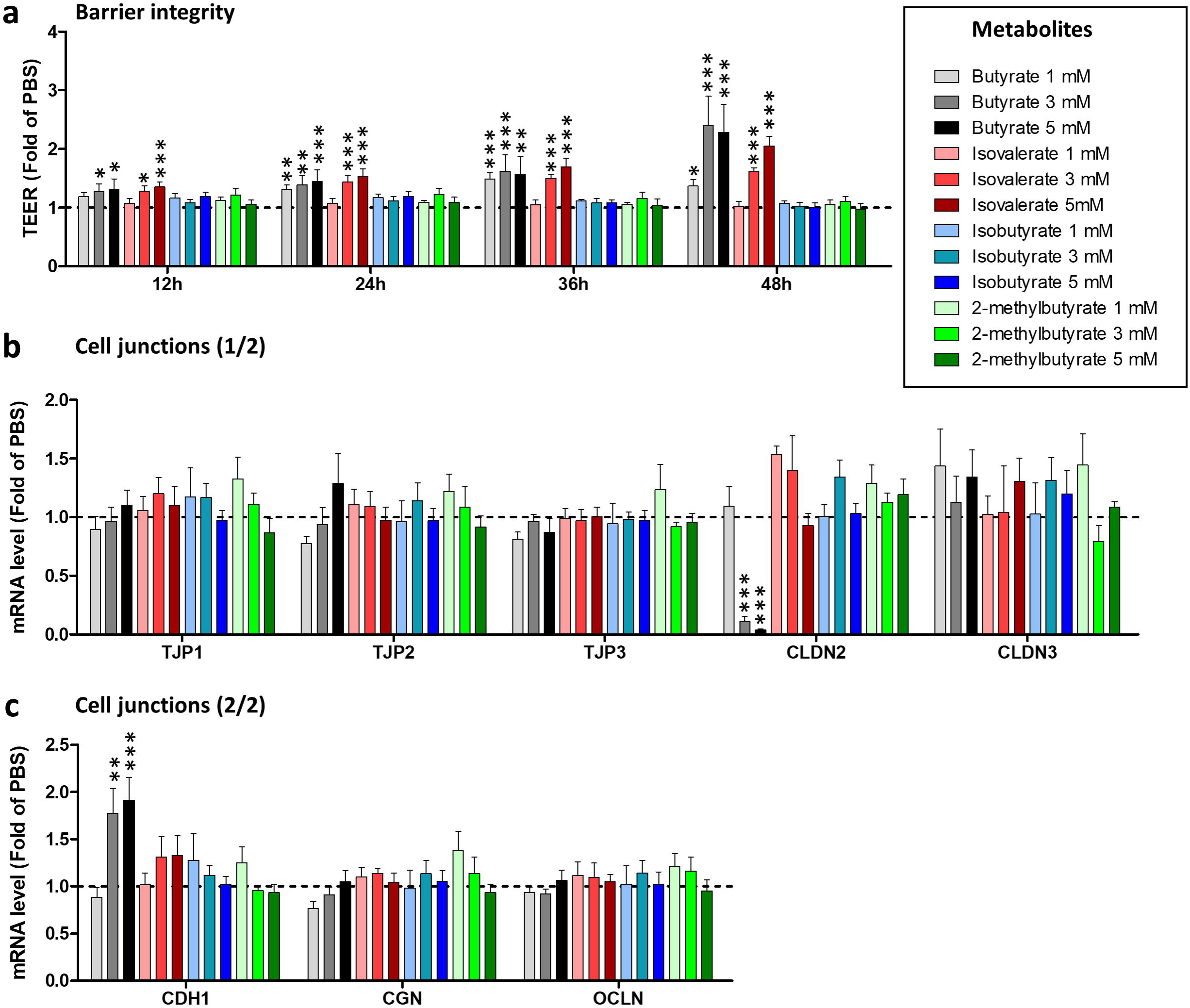
Effects of butyrate and branched-chain fatty acids on the epithelial barrier function. Cell monolayers derived from porcine ileum organoids were treated for 48h with bacterial metabolites (butyrate, isovalerate, isobutyrate, 2-methylbutyrate) at 1 or 3 or 5 mM or with vehicle (PBS, negative control). a) Transepithelial electrical resistance (TEER) according to time shown as means and SEM, n>5/group. Data are expressed relative to the control condition (PBS), represented by the dotted line (y=1). b - c) Relative expression of genes coding for cell junction proteins are represented as means and SEM, n>3/group. Data are expressed relative to the control condition (PBS), represented by the dotted line (y=1). *: P<0.05 versus control group, **: P<0.01 versus control group, ***: P<0.001 versus control group.

The antimicrobial protein SLPI was strongly increased by butyrate (1, 3 and 5 mM) and isovalerate (3 and 5 mM) (Figure 5A). Butyrate (5 mM) increased the expression of the microbial sensor TLR2 (Figure 5B). Conversely, butyrate (3 and 5 mM) decreased the expression of the TLR4, the lipopolysaccharide binding protein LBP, the TLR adaptor protein MYD88 and BAX, whereas it increased the expression of NFKB2 (Figures 5B and C). Butyrate (1, 3, and 5 mM) strongly increased the expression of the cytokine CXCL8, which was also upregulated by isovalerate (3 and 5 mM) and isobutyrate (1, 3 and 5 mM) (Figure 5D). Butyrate (3 and 5 mM) also upregulated the gene expression of the cytokine CCL20 (Figure 5D). Isovalerate (3 and 5 mM) was the only metabolite that increased the gene expression of the antioxidant enzymes GPX1 and SOD1 (Figure 5E). Conversely, butyrate (3 and 5 mM) decreased the expression of GPX2 and the highest concentration of butyrate (5 mM) also decreased the expression of the DUOX2 and SOD1 (Figure 5E).

**Figure 5:**
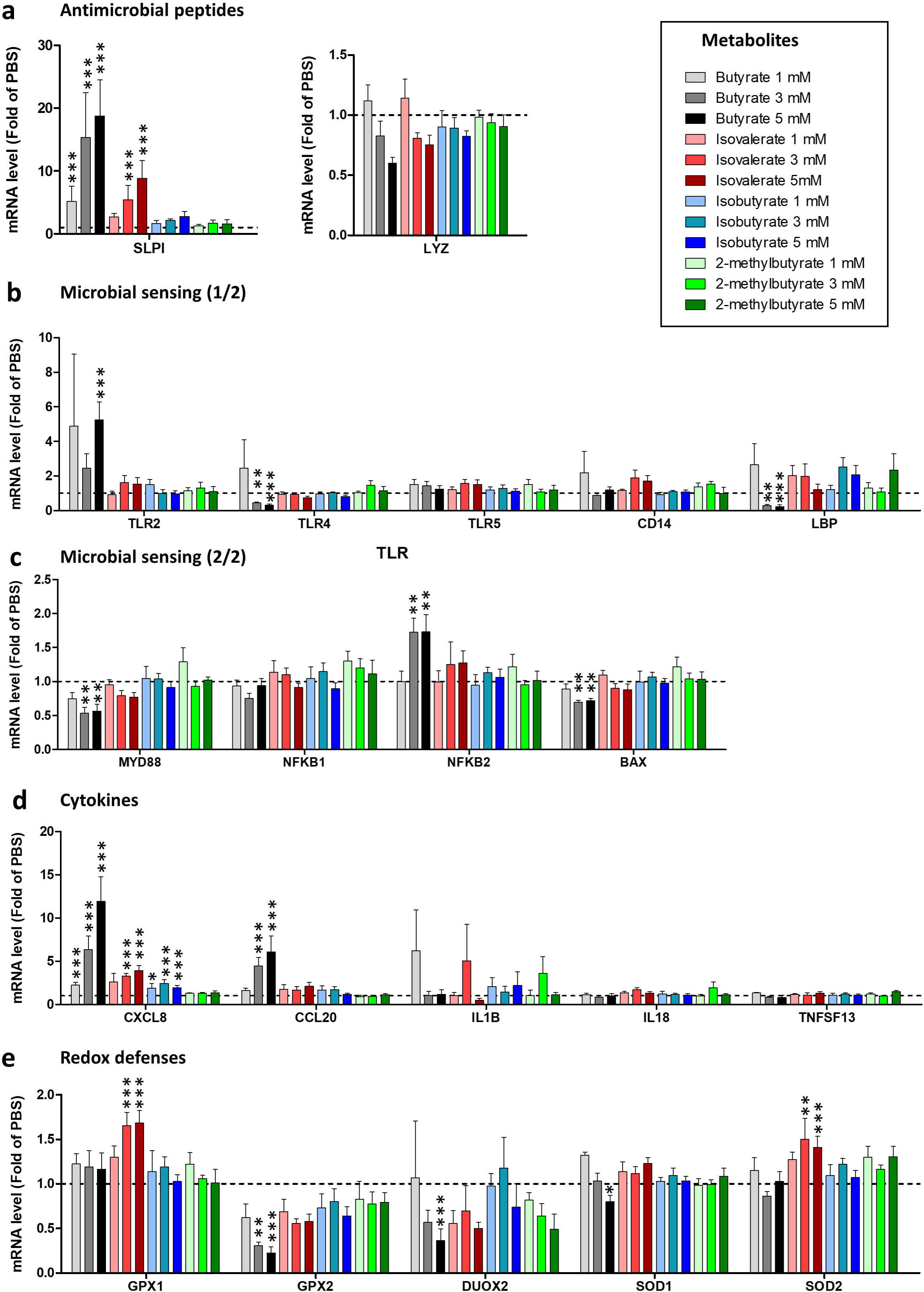
Effects of butyrate and branched-chain fatty acids on epithelial antimicrobial defenses. Cell monolayers derived from porcine ileum organoids were treated for 48h with bacterial metabolites (butyrate, isovalerate, isobutyrate, 2-methylbutyrate) at 1 or 3 or 5 mM or with vehicle (PBS, negative control). a - e) Relative gene expressions are represented as means and SEM, n>3/group. Data are expressed relative to the control condition (PBS), represented by the dotted line (y=1). *: P<0.05 versus control group, **: P<0.01 versus control group, ***: P<0.001 versus control group. a) Genes encoding antimicrobial peptides. b - c) Genes encoding proteins involved in microbial sensing. d) Genes encoding cytokines. e) Genes encoding proteins involved in redox defenses.

Butyrate (3 and 5 mM) strongly reduced the expression of marker genes for stem cells (OLFM4 and SMOC2) and progenitor cells (MKI67 and PCNA), while the expression of CFTR was reduced only at 5 mM concentration (Figure 6A). Isovalerate (5 mM) also reduced the expression of OFLM4. Conversely, the progenitor cell marker SOX9 was upregulated by the two highest concentration of butyrate (Figure 6B). Butyrate reduced the expression of absorptive cell marker genes (KRT20 and CA2) and of the glucose transporter SLC5A1 (Figure 6C). The highest concentration of butyrate (5 mM) reduced the expression of FABP1 while lower concentrations (1 and 3 mM) increased the expression of FABP6 (Figure 6D). Isovalerate (5 mM) also upregulated FABP6. Butyrate (3 and 5 mM) reduced the expression of the bile acid receptor encoded by the NR1H4 gene (Figure 6D). Butyrate (1, 3 and 5 mM) and isovalerate (3 and 5 mM) increased the expression of the peptide transporter PEPT1 (Figure 6E). Butyrate (3 mM) reduced the expression of the amino acid transporter SLC7A11 (Figure 6E). Butyrate (3 and 5 mM) and isovalerate (5 mM) increased the expression of ARG2 (Figure 6E).

**Figure 6:**
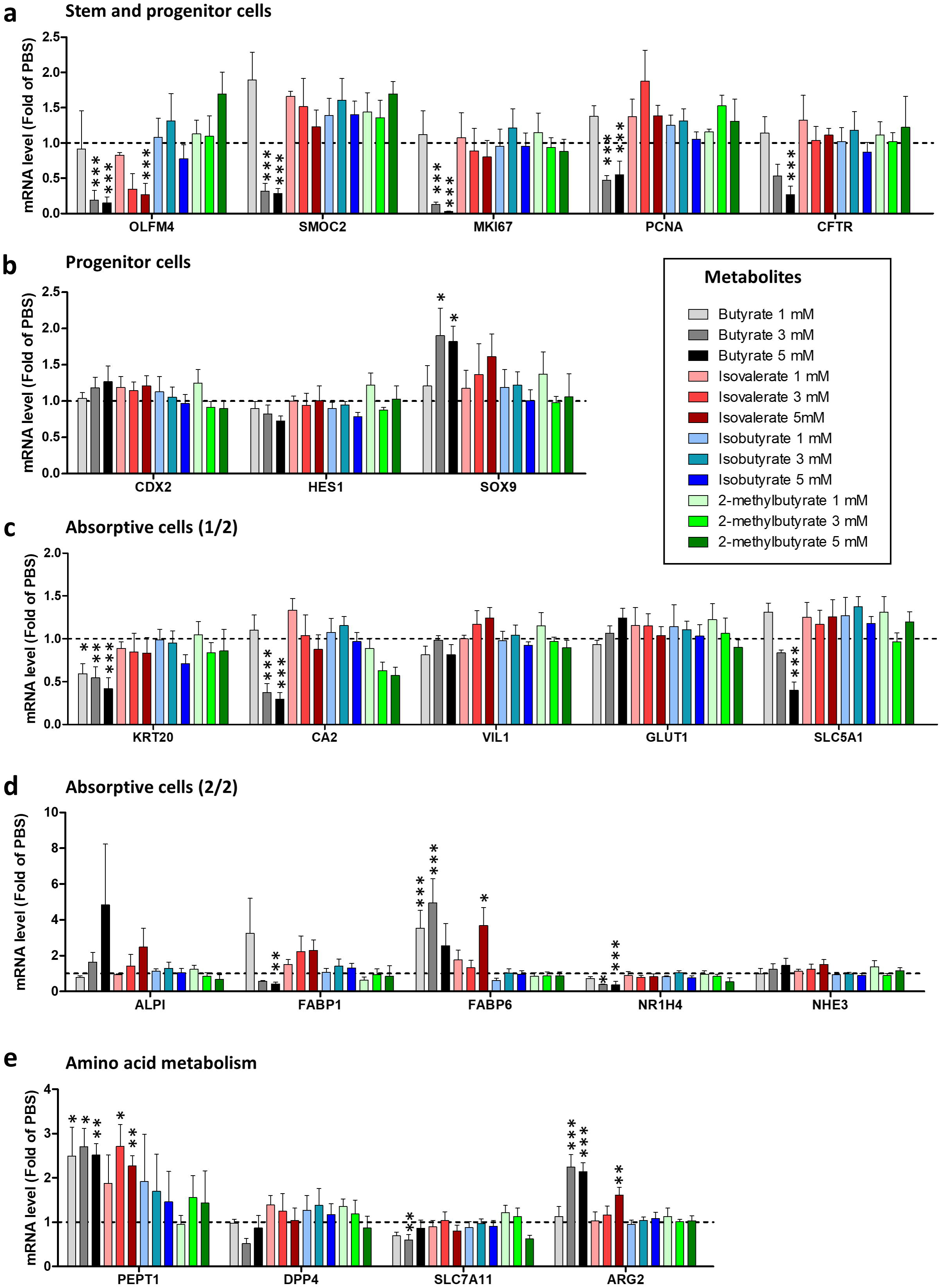
Effects of butyrate and branched-chain fatty acids on epithelial cell proliferation and absorptive lineage differentiation. Cell monolayers derived from porcine ileum organoids were treated for 48h with bacterial metabolites (butyrate, isovalerate, isobutyrate, 2-methylbutyrate) at 1 or 3 or 5 mM or with vehicle (PBS, negative control). a - e) Relative gene expressions are represented as means and SEM, n>3/group. Data are expressed relative to the control condition (PBS), represented by the dotted line (y=1). *: P<0.05 versus control group, **: P<0.01 versus control group, ***: P<0.001 versus control group. a - b) Marker genes for stem and progenitor cells. c - d) Marker genes for absorptive cells. e) Genes encoding proteins involved in amino acid metabolism.

Butyrate (3 and 5 mM) and isovalerate (5 mM) increased the gene expression of the enteroendocrine cell marker CHGA (Figure 7A). On the contrary, butyrate decreased the expression of the genes encoding the enterohormones glucagon (GCG) and peptide YY (PYY). The gene expression of the transmembrane mucin MUC1 was reduced by butyrate (1, 3 and 5 mM), isovalerate (3 and 5 mM) and 2-methylbutyrate at 5 mM (Figure 7B). Butyrate (3 and 5 mM) reduced the gene expression of the goblet cell markers ATOH1, TFF3 and MUC2 (Figure 7C). The highest concentration of isovalerate (5 mM) also reduced the expression of ATOH1, while the expression of TFF3 was reduced at both 3 and 5 mM (Figure 7C). Isobutyrate (5 mM) also reduced the expression of TFF3. Taken together, the dose-response experiment showed that isovalerate, in contrast to other BCFAs, improves epithelial barrier integrity and regulates the expression of genes involved in diverse epithelial functions.

**Figure 7:**
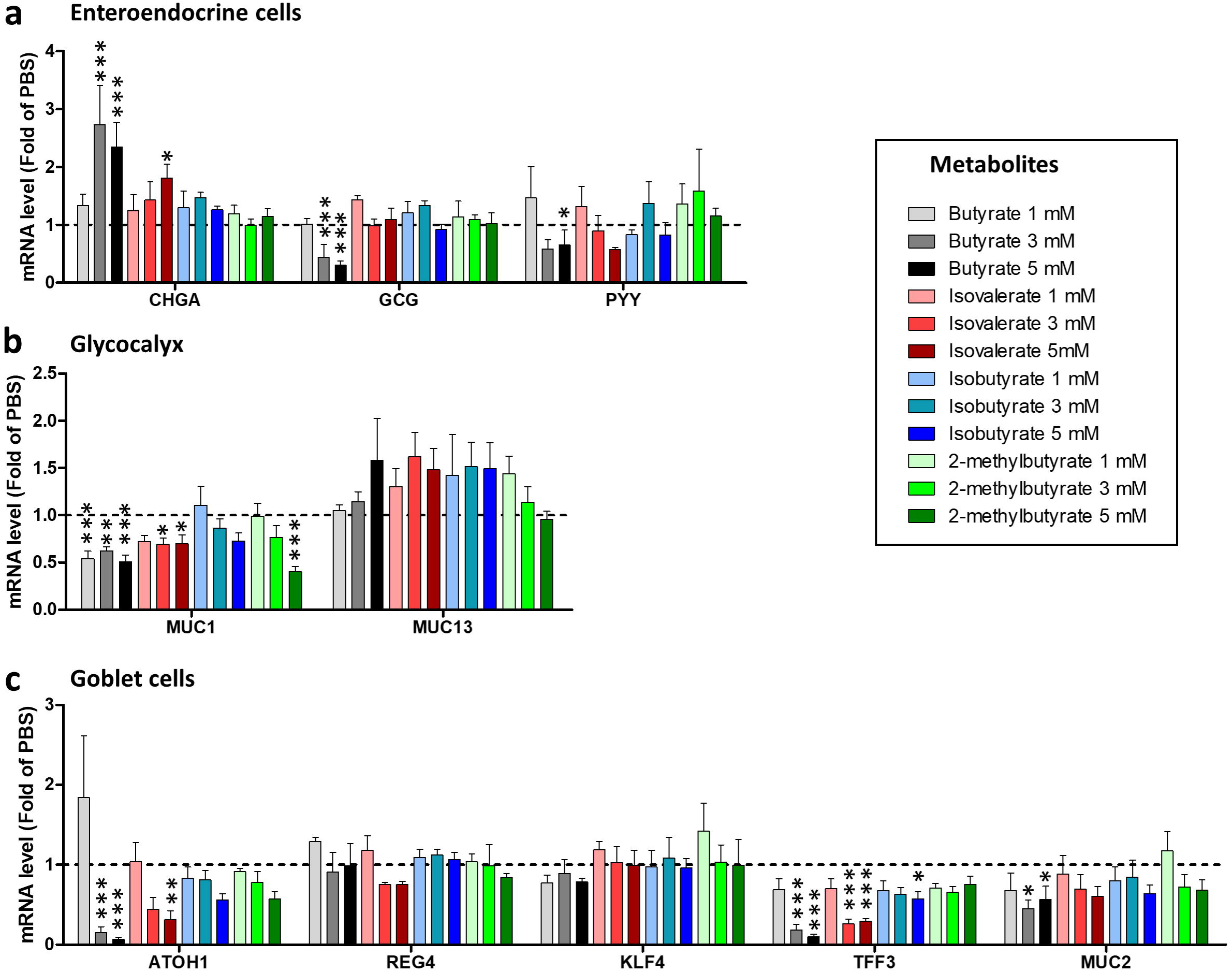
Effects of butyrate and branched-chain fatty acids on epithelial secretory cell differentiation and glycocalyx formation. Cell monolayers derived from porcine ileum organoids were treated for 48h with bacterial metabolites (butyrate, isovalerate, isobutyrate, 2-methylbutyrate) at 1 or 3 or 5 mM or with vehicle (PBS, negative control). a - c) Relative gene expressions are represented as means and SEM, n>3/group. Data are expressed relative to the control condition (PBS), represented by the dotted line (y=1). *: P<0.05 versus control group, **: P<0.01 versus control group, ***: P<0.001 versus control group. a) Marker genes for enteroendocrine cells. b) Genes encoding transmembrane mucins. c) Marker genes for goblet cells.

### Butyrate and branched-chain fatty acids are transported across cell monolayers derived from porcine ileum organoids

As a next step, we investigated whether the specific barrier promoting effects of isovalerate among BCFAs was linked to its transepithelial transport. To this aim, we analyzed by NMR the apical and basal culture media of the cell monolayers derived from porcine ileum organoids treated at the apical side with butyrate or BCFAs at 5 mM for 48h. Butyrate and all three BCFA were detected both in the apical and basolateral culture media at the end of the experiment (Figures 8A and B). These results indicate that butyrate and BCFAs are transported across the intestinal epithelium *in vitro* and exclude out the possibility that effects of isovalerate on the intestinal epithelium are related to a specific transepithelial transport.

**Figure 8:**
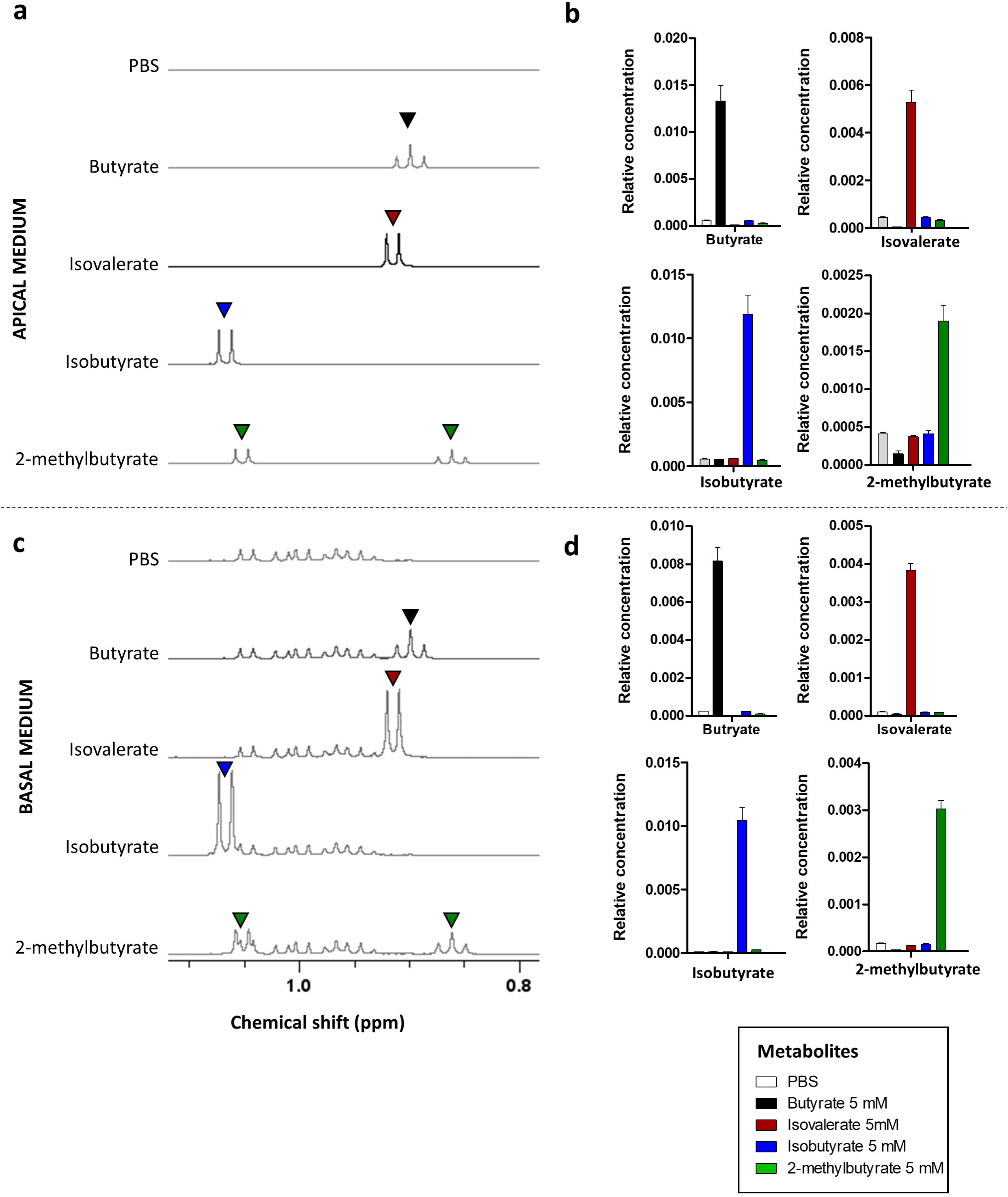
Transepithelial transport of butyrate and branched-chain fatty acids in cell monolayers derived from porcine ileum organoids. Cell monolayers derived from porcine ileum organoids were treated for 48h with bacterial metabolites (butyrate, isovalerate, isobutyrate, 2-methylbutyrate) at 5 mM or with vehicle (PBS, negative control). The metabolome of the apical and basal media collected at the end of the experiment were analyzed by nuclear magnetic resonance (NMR) spectroscopy. a, c) Representative NMR spectra of the apical (a) and basal (c) media in each experimental condition. Arrowheads indicate peaks corresponding to butyrate (black), isovalerate (red), isobutyrate (blue) and 2-methylbutyrate (green). b, d) Relative concentration of branched-chain fatty acids in the apical (b) and basal (d) media represented as means and SEM, n>4/group.

### Inhibition of histone deacetylases replicates the barrier promoting effects of butyrate and isovalerate

Histone deacetylase (HDAC) inhibition is a central mechanism by which short-chain fatty acids regulate gene expression in host cells (McBain et al. 1997). To assess the involvement of this mechanism in our model, we treated cell monolayers derived from porcine ileum organoids for 48h with butyrate (5mM) or isovalerate (5 mM) or TSA (1 µM), an HDAC inhibitor structurally dissimilar to short-chain fatty acids. TSA mimicked the protective effects of butyrate and isovalerate on epithelial barrier integrity, as indicated by the increased TEER from 24h onwards (Figure 9A). Butyrate, isovalerate and TSA also reduced the paracellular permeability to FITC-dextran 4 kDa (Figure 9B). In addition, TSA strongly upregulated SLPI gene expression, reflecting the effects of butyrate and, to a lesser extent, isovalerate (Figure 9C). Taken together, these results indicate that inhibition of HDAC replicates the effects of butyrate and isovalerate on cell monolayers derived from porcine ileum organoids.

**Figure 9:**
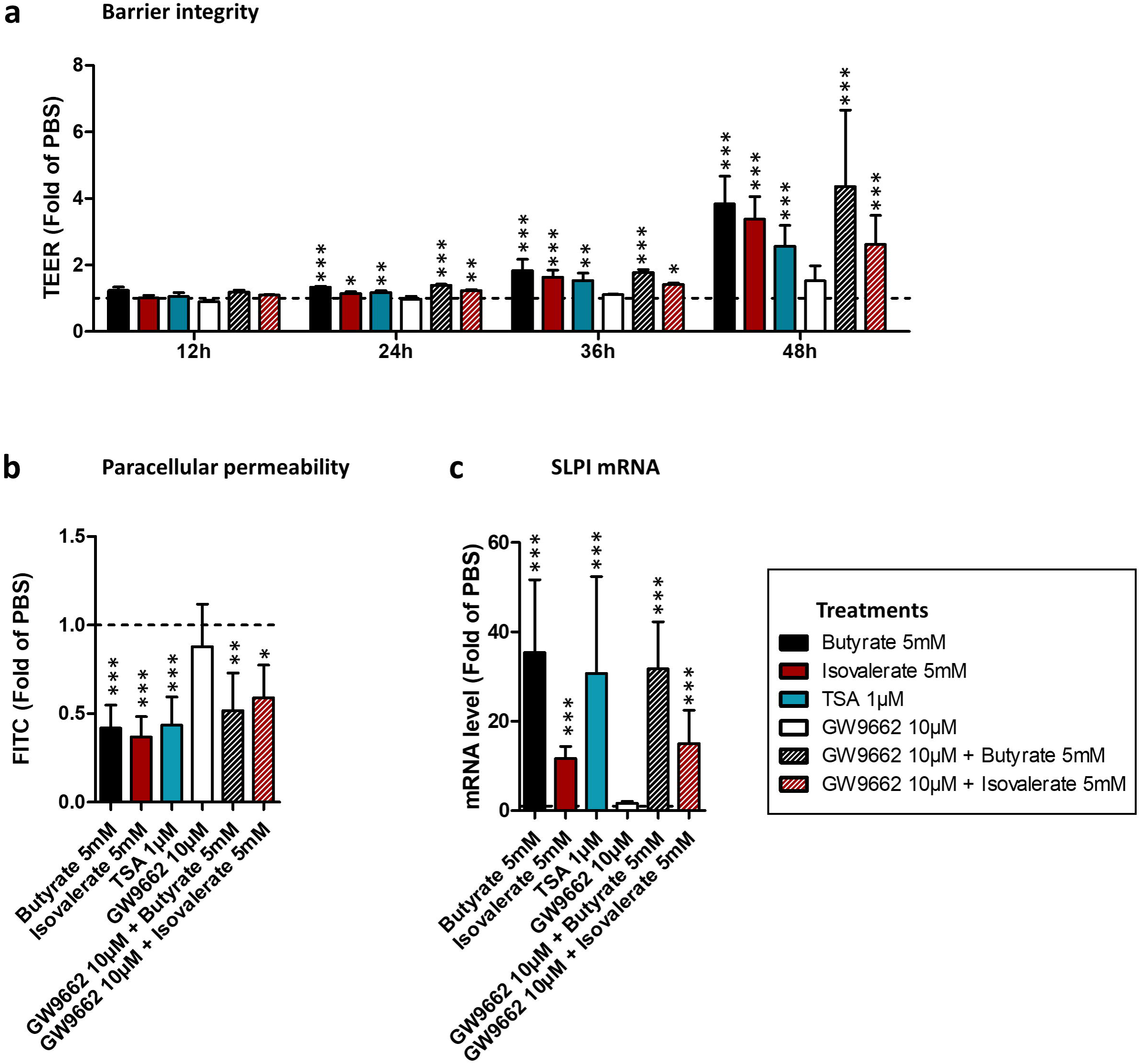
Mode of action of butyrate and isovalerate on the epithelial barrier function. Cell monolayers derived from porcine ileum organoids were treated for 48h with bacterial metabolites (butyrate, isovalerate) at 5 mM combined or not with the PPARγ inhibitor GW9662 10µM or Trichostatin A (TSA) 1µM or with vehicle (PBS, negative control). a) Transepithelial electrical resistance (TEER) according to time shown as means and SEM, n>3/group. b) Paracellular permeability evaluated by the transport of FITC-dextran 4 kDa shown as means and SEM, n>3/group. c) SLPI gene expression shown as means and SEM, n>3/group. a - c) Data are expressed relative to the control condition (PBS), represented by the dotted line (y=1). *: P<0.05 versus control group, **: P<0.01 versus control group, ***: P<0.001 versus control group.

Activation of peroxisome proliferator-activated receptor γ (PPARγ) is another mechanism underlying the effects of short-chain fatty acids on host cells (Wang et al. 2023). To investigate if PPARγ activation mediates the effects of butyrate and isovalerate in our model, we treated cell monolayers derived from porcine ileum organoids with butyrate (5 mM) or isovalerate (5 mM) in presence of the PPARγ inhibitor GW9662 (10 µM). Our results show that GW9662 had no effect on TEER, FITC-dextran 4 kDa permeability and SLPI gene expression (Figures 9A-C). The effects of butyrate and isovalerate on epithelial cells were similar with or without GW9662. These data suggested that butyrate and isovalerate improved the epithelial barrier function independently of PPARγ signaling.

### Bacterial BCFA production through a non-canonical pathway

Following our identification of the beneficial effects of isovalerate on intestinal barrier function *in vitro*, we wanted to identify which bacteria in the porcine gut microbiota could produce this metabolite. The biosynthesis of BCFAs involves two main steps before the conversion into the final product (Kaneda 1977). Precursor branched-chain amino acids are first deaminated into their α-ketoacid forms through the transamination of glutamate, and then these are oxidatively decarboxylated to the corresponding CoA derivatives, before being converted into BCFAs (Figure 10A). While the first reaction is carried out by a branched-chain amino acid aminotransferase (BCAT), different enzymes have been described to perform the second step. In bacteria such as *Bacillus subtilis* and *Streptomyces avermitilis* this reaction is carried out by the branched-chain α-ketoacid dehydrogenase complex (BCKDH), a large multienzyme complex comprising several subunits (Wang et al. 1993; Denoya et al. 1995). Recently, an alternative BCFA pathway was proposed in the gut commensal bacteria *Clostridium sporogenes* involving PorA, described as a member of the pyruvate:ferredoxin oxidoreductase (POR) superfamily (Guo et al. 2019). Like BCKDH, PorA is CoA-dependent and contains thiamine diphosphate, but, in contrast with BCKDH, carries iron-sulfur clusters and requires a ferredoxin as electron donor instead of nicotinamide adenine dinucleotide (Heider et al. 1996).

**Figure 10:**
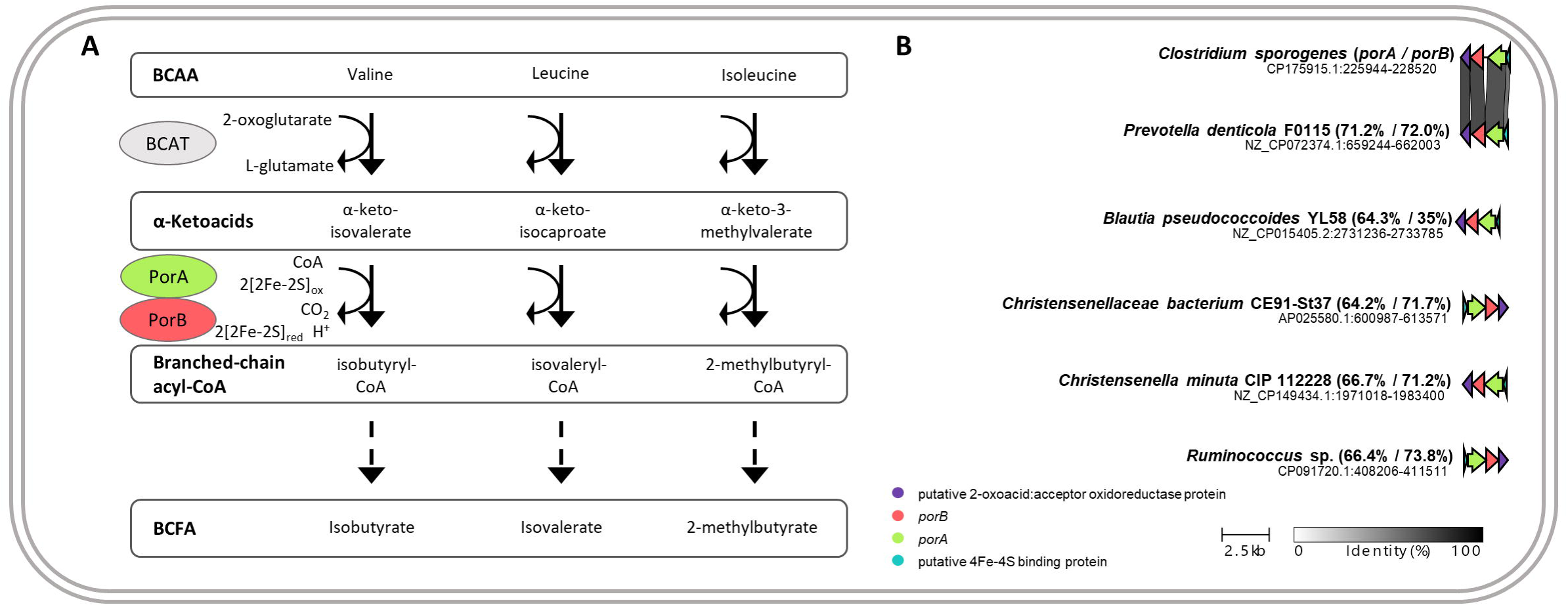
Branched-chain fatty acid metabolism in bacterial members of the pig gut microbiota. a) Predicted branched-chain fatty acids (BCFA) metabolic pathway involving a branched-chain amino acid (BCAA) aminotransferase (BCAT), a pyruvate:ferredoxin oxidoreductase (PorA) and a thiamine pyrophosphate-dependent enzyme (PorB). b) Distribution of the *porA* locus in species of prevalent genera of the pig gut microbiota. *porA* and *porB* homologous gene identity compared to *C. sporogenes*. Accession numbers as well as the exact position of the locus is indicated for each strain.

We performed *porA* homology searches on genera previously identified as abundant in the pig gut microbiota using cblaster (Gilchrist and Chooi 2021; Alberge et al. 2024). The results revealed that *porA* was found in some species with a high degree of similarity ranging from 64 to 71% identity, namely in *Prevotella*, *Blautia*, *Chirstensenella* and *Ruminococcus* species (Figure 10B). Interestingly, *porA* was consistently clustered with a thiamine pyrophosphate-dependent enzyme encoding gene, *porB* (between 35 and 73% identity), as well as a putative 4Fe-4S-binding protein and a putative 2-oxoacid:acceptor oxidoreductase (Figure 10B). On the contrary, no homologues of these genes were identified in species from other dominant genera of the porcine gut microbiota such as *Terrisporobacter*, *Faecalibacterium*, *Lactobacillus* and *Limosilactobacillus* (data not shown).

## Discussion

Our screening of bacterial metabolites on organoid cell monolayers derived from the porcine ileum showed that butyrate was the metabolite with the greatest effect on epithelial cells. Indeed, we found that butyrate enhanced the epithelial barrier function, regulated the expression of tight junction proteins and innate immune defense systems, reduced cell proliferation and goblet cell differentiation while promoting the differentiation of absorptive and enteroendocrine cells. These results are consistent with previous studies (Peng et al. 2009; Yan and Ajuwon 2017; Pearce et al. 2020; Beaumont et al. 2023; Alberge et al. 2024) and confirm that our experimental system is suitable to identify bacterial metabolites able to regulate epithelial functions. Accordingly, we identified several genes whose expression was regulated by isovalerate (SLPI, FABP6, CHGA) and tryptamine (BAX, FABP6, ARG2), when tested at 1 mM. Protective effects of the tryptophan-derived metabolite tryptamine on the intestinal epithelium were described previously (Williams et al. 2014; Bhattarai et al. 2018, 2020) and we decided to focus our study on isovalerate and other BCFAs, whose concentration can reach up to 5 mM in the human and pig intestine (Cummings et al. 1987; Macfarlane et al. 1992; Beaumont et al. 2017; Rios-Covian et al. 2020; Wang et al. 2023; Alberge et al. 2024). Interestingly, we found that isovalerate tested at 3 or 5 mM regulated the expression of more genes when compared to 1 mM, while the two other BCFAs only regulated the expression of one gene at the highest concentration tested (CXCL8 by isobutyrate and MUC1 by 2-methylbutyrate). These results suggest that the effects of isovalerate were dose-dependent and specific to the structure of the carbon chain of this BCFA.

The effects of isovalerate on epithelial cells were in most cases also observed with butyrate, which suggest a common mode of action. Indeed, both isovalerate and butyrate enhanced the barrier function as indicated by TEER measurement, upregulated the expression of genes involved in innate immunity (SLPI, CXCL8), and markers of absorptive cells (FABP6, PEPT1, ARG2) and enteroendocrine cells (CHGA) while reducing the expression of the stem cell marker OLFM4 and of mucus related genes (MUC1, ATOH1, TFF3). The common mode of action of butyrate and isovalerate may involve inhibition of HDAC. Indeed, short-chain fatty acids able to inhibit HDAC were previously shown to be 3-5 carbon in length and lack substitution at the 2-position (McBain et al. 1997), which corresponds to the molecular structure of both butyrate and isovalerate but not of isobutyrate and 2-methylbutyrate that had no effect on the epithelium in our study. Furthermore, 2-methylbutyrate has the same mass as isovalerate and no effect is observed for this metabolite in the studied conditions, reinforcing the notion that the structure of BCFAs mediates their biological effects. The potential role of HDAC inhibition mediating the effects of isovalerate and butyrate in our experimental model is further supported by our experiments showing that the structurally unrelated HDAC inhibitor TSA also enhanced the barrier function, reduced paracellular permeability and induced the expression of SLPI, as observed with both butyrate and isovalerate.

We also found that isovalerate upregulated the gene expression of the antioxidant enzymes GPX1 and SOD2, which was not observed with butyrate. This isovalerate-specific effect may thus involve additional mechanisms not shared with butyrate. A study showed the post-translation modification SUMOylation was increased by BCFAs in human intestinal cell lines through transient induction of reactive-oxygen species (ROS) (Ezzine et al. 2022). The induction of ROS by isovalerate could therefore explain the increased gene expression of enzymes involved in antioxidant responses that we observed in porcine organoid cell monolayers. Another study also indicated that some of the biological effects of isovalerate on host cells are mediated by PPARγ signaling (Wang et al. 2023). However, we found that the PPARγ inhibitor GW9662 did not prevent the effect of isovalerate on the barrier function or SLPI induction in cell monolayers derived from porcine ileum organoids. Other potential modes of action of isovalerate identified previously and involving the G protein-coupled receptors Olfr558 or the mammalian target of rapamycin (mTOR) signaling remain to be evaluated in the context of the regulation of epithelial barrier function (Bellono et al. 2017; Choi et al. 2021; Wang et al. 2023).

Our results showing that isovalerate can cross the epithelial barrier *in vitro* suggest that this bacterial metabolite may have biological effects behind the epithelium and notably on immune cells localized in the intestinal *lamina propria*. Indeed, several studies indicated that isovalerate modulates immunoglobulin-A secreting plasma cells (Guo et al. 2019; Wang et al. 2023), cytokine expression in Th17 and B cells and macrophage polarization (Wang et al. 2023). Additionally, BCFAs isovalerate and isobutyrate can be sensed by bacteria, which can modulate their by pathogenicity through transcriptional regulations, as demonstrated for *Campylobacter jejuni* (Ruiz et al. 2024). BCFAs may therefore play an important role in bacteria-pathogen and bacteria-host communication in the gut.

Modulating the production of isovalerate by the microbiota therefore appears as a potential strategy to promote gut health. BCFA production was demonstrated *in vitro* in several gut bacteria including *Clostridium* species*, Bacteroides* species*, Parabacteroides merdae,* and *Megasphaera elsdenii* (Allison 1978; Elsden and Hilton 1978; Qiao et al. 2022; Wang et al. 2024). Genetic engineering experiments demonstrated that the *porA* gene is necessary for the production of BCFA in *Clostridium sporogenes* and *Parabacteroides merdae* (Guo et al. 2019; Qiao et al. 2022). In humans, BCFA production has been mainly attributed to *Clostridium* and *Bacteroides* strains (Rios-Covian et al. 2020). Instead, our *in silico* analysis indicated that other bacteria such as species from *Prevotella*, *Blautia*, *Christensenella* and *Ruminococcus*, which are highly prevalent in the pig gut microbiota (Alberge et al. 2024), could be BCFA producers in pigs, as suggested by the presence of *porA* homologues in their genomes. The conserved genetic organization observed between *porA* and thiamine pyrophosphate-dependent enzyme and 4Fe-4S-binding protein encoding genes would provide a plausible electron transfer for a viable BCFA production pathway, although further culture-based studies will be necessary to confirm this. Colonization of mice with *Clostridium sporogenes* or *Parabacteroides merdae* induces the accumulation of BCFA in the gut (Guo et al. 2019; Qiao et al. 2022; Krause et al. 2024), which suggests that such bacterial species could be used as isovalerate-producing probiotics to improve gut health. Nutritional strategies could also be used to promote the production of isovalerate by gut bacteria through increasing the supply of leucine, the amino acid precursor of isovalerate. The simplest approach would be to increase the level of protein intake which was shown previously to increase the concentration of BCFAs, including isovalerate (Pieper et al. 2012; Beaumont et al. 2017). However, high-protein diets also increase the production of other amino acid derived bacterial metabolites with potential detrimental effects such as p-cresol and hydrogen sulfide (Andriamihaja et al. 2015; Beaumont et al. 2016), and a high protein intake is suspected to be associated with digestive disorders (Gilbert et al. 2018). An alternative would be to use leucine supplementation to specifically bring the precursor of isovalerate to the gut microbiota. In order to delay the rapid absorption of free leucine in the upper small intestine, the utilization of protected-leucine (e.g., coated in a lipid matrix) may be a promising strategy to release leucine to the gut microbiota in the distal intestine for the production of isovalerate (Beaumont et al. 2022).

In conclusion, our study shows that the leucine-derived gut microbiota BCFA isovalerate improves the epithelial barrier function *in vitro.* This result suggests that targeting the bacterial production of isovalerate may be a promising strategy to promote gut health. The identification of species from *Prevotella*, *Blautia*, *Christensenella* and *Ruminococcus* as potential producers of BCFAs in the pig gut microbiota paves the way for the development of probiotics selected based on their metabolic capacity.

## ACKNOWLEDGMENTS

The authors thank the GenoToul platform for metabolomics (MetaToul-Axiom, Toulouse, France) and the genomic platform GENTYANE (INRAE, Clermont Ferrand, France).

## AUTHOR CONTRIBUTIONS

MB and TCD designed research; MB, CL and EJ conducted research; MB and CMV analyzed data; MB wrote the initial draft; CMV, CL, EJ, SL and TCD reviewed the manuscript. All authors approved the final manuscript.

## DATA AVAILABILITY STATEMENT

All data are contained in the article and the supporting information.

